# Integrative proteo-genomic profiling uncovers key biomarkers of lapatinib resistance in HER2-positive breast cancer

**DOI:** 10.1101/2024.11.08.621581

**Authors:** J Steggall, V Rajeeve, N Al-Subaie, A Hayat

## Abstract

Drug resistance is a major obstacle to the long-term effectiveness of cancer therapies. Approximately 70% of breast cancer patients relapse after 5 years of treatment, and the lack of biomarkers associated with drug resistance translates to poor prognosis in clinic. Previous research has utilised omics approaches to uncover biomarkers driving drug resistance, with a strong emphasis on genetic mutations. Here, we identified a nine-marker signature associated with resistance to lapatinib in a HER2-positive breast cancer model using a target discovery approach by employing an integrative multi-omics strategy, combining ATAC-seq, RNA-seq, and proteomics. We found that 7 markers in the drug resistance-signature had not been previously found to be implicated in HER2 positive breast cancer. We counterintuitively found that drug resistant cells have restrictive chromatin accessibility with reduced gene expression associated with limited total proteome changes. However, upon closer look, we identified that the drug resistance-signature had increased chromatin accessibility closer to the transcriptional start sites of those genes and are highly differentially expressed across the three datasets. Our data show that despite the overall transcriptional and proteomic landscape showing limited changes, there are several markers that are highly expressed, which correlate with increased anchorage-independent and invasive phenotype *in vitro* in lapatinib resistant cells compared to cancer cells. Our results demonstrate that disease aggressiveness can be related to reduced chromatin and gene expression dynamics. We anticipate that the resistant signature identified here using integrative target discovery approach can be applied to complex, representative models and validated before they can be targeted by suitable therapeutic agents.

## Introduction

Breast cancer is the most commonly diagnosed cancer worldwide, with 1 in 5 cases exhibiting the overexpression of the Human Epidermal Growth Factor Receptor 2 (HER2). Despite significant advancements in treatment in recent years, more than 70% of patients relapse after 5 years, indicating the need to interrogate the mechanisms of acquired drug resistance (1). Integrative multi-omics approaches are a powerful tool in aiding to elucidate the aberrant changes driving acquired drug resistance. Identifying novel biomarkers and understanding the molecular mechanisms contributing to acquired drug resistance are vital for tackling cancer dissemination and metastasis.

Lapatinib, a dual kinase inhibitor, targets HER2 and Epidermal Growth Factor Receptor (EGFR) domains, thereby preventing phosphorylation and subsequent signal transduction of the mitogen-activated protein kinase (MAPK) and phosphoinositide 3-kinase (PI3K)/Akt pathways, leading to cell death (2). Lapatinib is frequently used in combination with other therapeutic agents for treating high-risk metastatic breast cancer patients, particularly in cases where they display resistance to trastuzumab, a first line therapeutic agent for HER2 positive breast cancer (3,4). Despite initial efficacy, most patients inevitably develop resistance to lapatinib and, sadly, succumb to disease progression and mortality (5). A major challenge remains in how we overcome this resistance. The selective pressure as a result of targeted therapies on HER2 leads to rapid evolution of acquired drug resistance mechanisms, reducing the efficacy of personalised medicine approaches resulting in disease relapse (6).

A growing number of changes have been identified in the genome and the epigenome of breast cancer cell lines, with the upregulation of *SCIN* and *GIRK1* playing a key role in promoting proliferation and inhibiting apoptosis. Alongside this, the downregulation of *EGR1,* leads to the overexpression of multidrug resistance protein 1 *(MDR1)*, contributing to poor prognosis and therapeutic resistance (7–11). However, despite these advancements in identifying several proteins associated with acquired drug resistance, the transcriptome and particularly the chromatin dynamics remains largely unexplored, informing our integrative approach to perform deep-sequencing analysis of high HER2 expressing cancer cells (SKBR3) and its lapatinib-resistant counterpart (SKBR3-L) using multi-omics. Chromatin accessibility is a key regulator of gene expression and alterations in the chromatin state can lead to significant changes in transcriptional activity, which are eventually translated into proteins, the effectors of cell function (12). Nevertheless, DNA alterations and changes in RNA expression do not always translate into protein expression patterns that cause distinct biological changes in cell function at the level where targeted therapies work (13). Therefore, using a combination of omics techniques that allow us to characterise a snapshot of the molecular landscape at several different layers of regulations is likely to define molecular changes critical for acquired drug resistance.

After we identified specific genomic regions and transcription factors that play a crucial role in regulating gene expression using (assay of transposase-accessible chromatin using sequencing) ATAC-seq and RNA-seq. We conducted an unbiased global proteome analysis of the same cells to catalogue the intricate workings of lapatinib resistance at protein level. By exploring the genome, transcriptome, and proteome at each level, we developed a comprehensive understanding of drug resistance in these cells. This integrated approach helped uncover the underlying changes that may contribute to drug resistance. These findings could pave the way for further validation of biomarkers in additional models, which could lead to clinical validation. Ultimately, this could improve the efficacy of HER2-targeted therapies using a sequence of biomarker-guided therapies that result in a cure (14). We unexpectedly discovered a significant decrease in chromatin accessibility, which aligned with changes in gene expression patterns in the drug-resistant cells. These changes were associated with limited alterations in the proteome. However, certain regions of the genome remained highly accessible at chromatin level, near the promoter, which could be contributing to the aggressive nature of drug-resistant cells.

## Results

### Lapatinib resistance evokes aggressive *in vitro* transformation

To explore the stochastic changes associated with acquired drug resistance, we obtained SKBR3 cells, which express high levels of HER2 protein with an established cell line model of acquired resistance to lapatinib, SKBR3-L cells (Fig. 1A) (15). We first tested the robustness of the system by analysing the HER2 protein expression using immunofluorescence. HER2 protein was seen in SKBR3 and SKBR3-L cells but not in the negative control as expected (Fig. 1B; supplementary fig. 1A). We next sought to establish whether there were differences in the morphological profiles between the two cell types. Cells were embedded into a 3D matrix and phase-focus quantitative time-lapse imaging was performed over 15 hours, allowing for a detailed examination of cell behaviour and morphology. Intriguingly, automated tracking revealed initial subtle decreases in perimeter in SKBR3-L cells which then stabilised, in contrast to the more pronounced perimeter reduction in SKBR3 cells (Fig. 1C). Throughout the experiment, SKBR3-L cells exhibited consistently reduced sphericity, assuming more irregular shapes and protrusions indicative of invasive capabilities. Additionally, despite an overall reduction in size, SKBR3-L maintained a larger cellular area, suggesting not only structural but also potentially functional adaptations conducive to invasive growth and metastatic potential as cells attempt to colonise the remaining space. We carried out soft agar colony formation assays, a gold standard for assessing *in vitro* transformation to test whether drug-resistant cells have higher transformational potential compared to cancer cells. SKBR3-L cells consistently formed enlarged structures in soft agar compared to SKBR3 cells across three different sizes, underscoring their highly transformed and drug-resistant nature (Fig. 1D and 1E).

**Figure 1.**
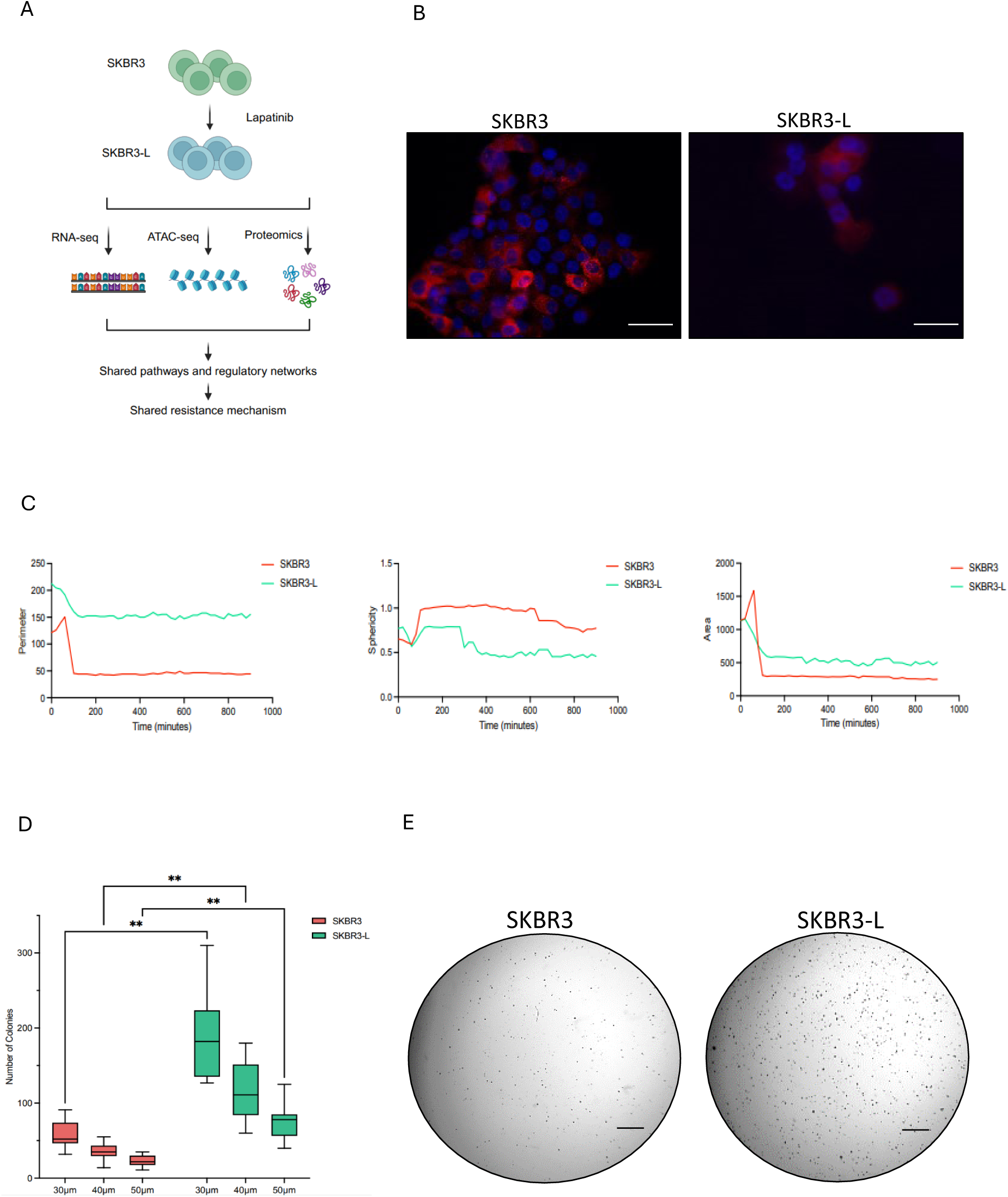
Phenotypic characteristics of SKBR3 and SKBR3-L cells. **(A)** Schematic diagram of the biological model we adapted, and the experimental approaches taken for the duration of the study. SKBR3 cells were made resistant to increasing concentrations of lapatinib. Various functional assays were used to characterise phenotypic changes before multi-omics analysis were performed to identify biomarkers in lapatinib acquired drug resistance. **(B)** Fluorescence microscopy confirmed the expression of HER2 protein in both cell types. 40x magnification. Scale bars: 100μm. **(C)** Cells were grown in 3D cell cultures and immediately imaged for 15 hours. Metrics such as area, sphericity and perimeter were automatically quantified and plotted as line graphs. **(D and E)** Anchorage-independent growth of cells was measured over 3 weeks after cells were grown in 0.35% ultra-pure agarose. Representative microscopic images of colonies stained with Crystal Violet are shown. Three biological replicates expressed as mean ± SEM. Analysis performed using two-way ANOVA followed by Holm-Šídák post hoc test; **p<0.005. 4x magnification. Scale bars: 100μm.

### Identification of distinct reduction in transcriptomic profile in acquired drug resistance

To characterise how the genomic landscape differs between cancer cells and their lapatinib resistant counterparts, we performed RNA-seq to determine gene expression patterns between the two cell types. The transcriptional programme of breast cancer cells and drug-acquired resistant cells differed significantly. Principal component analysis (PCA) separated the samples into two distinct clusters, confirming the difference between sensitive and resistant cell types (Fig. 2A). Notably, after the development of resistance, the cell line adopted a similar genetic profile, suggesting the emergence of a common adaptive response to therapy.

**Figure 2.**
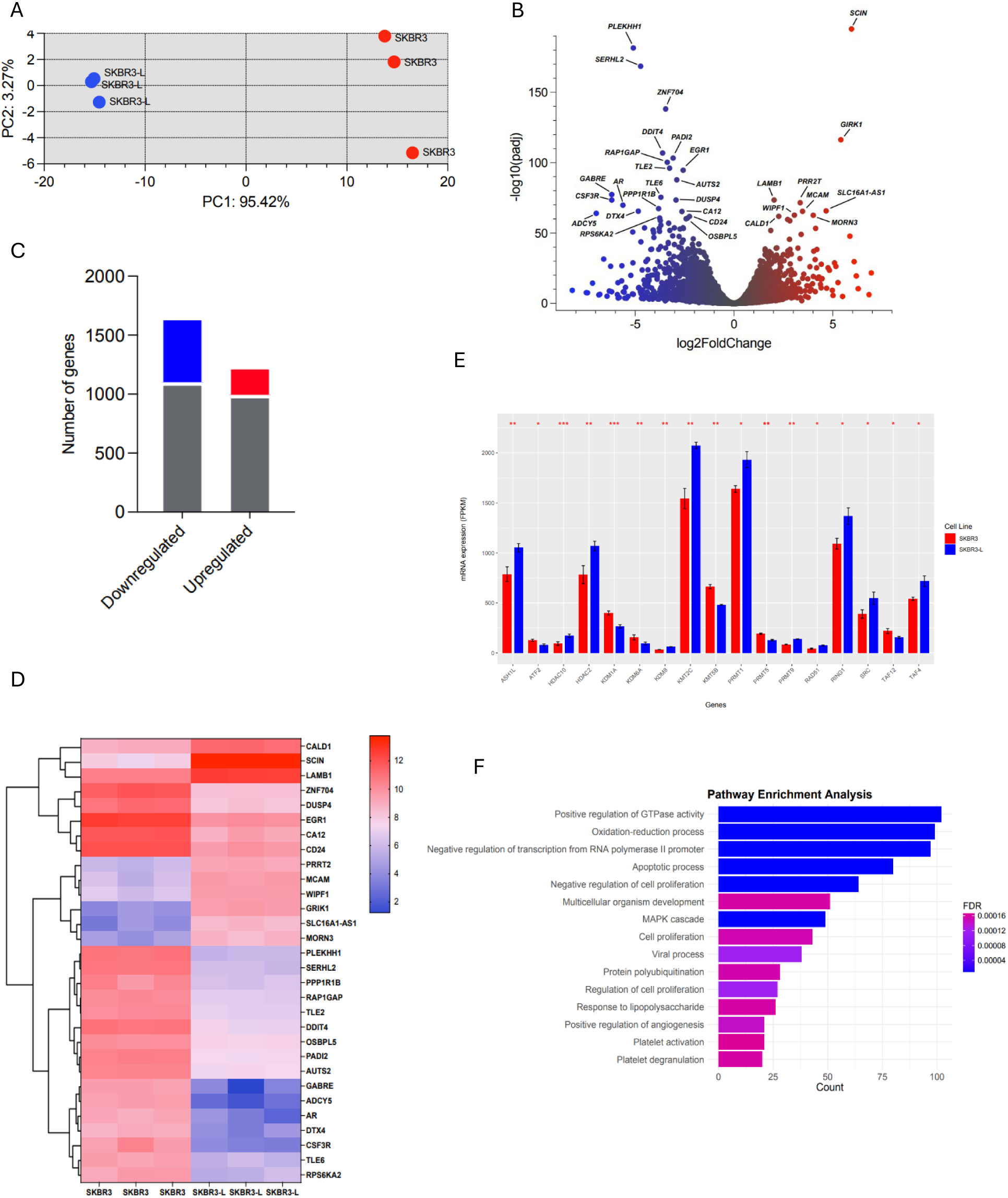
Differential gene expression between SKBR3 and SKBR3-L cells. **(A)** PCA plot displaying variance in gene expression profiles between SKBR3 (red) and SKBR3-L (blue). **(B)** Number of differentially expressed genes shown by the volcano plot in SKBR3 compared with SKBR3-L cells, downregulated genes are shown in blue with log2fold change < -2, adjusted p value < 0.05. Upregulated genes are shown in red with log2fold change > 2, adjusted p value < 0.05. The top 30 significantly differentially expressed are genes labelled. **(D)** Heatmap of the top 30 significantly differentially expressed genes are shown per individual replicate. **(E)** Gene expression of the significant epigenetic regulatory genes between SKBR3 and SKBR3-L cells. **(F)** Biological pathways associated with significant differentially expressed genes. For all this analysis, three biological replicates are expressed as mean ± SEM; *p<0.05, **p<0.005, ***p<0.0005.

We assessed the differential gene expression between these two groups and discovered that 8.5% were highly expressed genes with log2 fold > 1, p value < 0.05. Conversely, ∼19% of the genes were downregulated with log2 fold < -1, p value < 0.05 (Fig. 2B and 2C). We confirmed the differential upregulation of previously known genes in such as *SCIN, CALD1,* and *MCAM*, which have been associated with aggressive, basal-like phenotype, and in acquired drug resistance in cancer (16–18). However, we found novel genes such as *MORN3* with an unknown function, and *WIPF1*, which is an important differentiation marker but is not known to be involved in acquired drug resistance, both of which could be important as potential therapeutic resistance targets. Intriguingly, we identified *EGR1* to be significantly downregulated, which is implicated in breast cancer resistance and an important epigenetic regulator in methylation context, but whose function in HER2 positive breast cancer has yet to be understood (19). *CD24*, a critical breast stem marker, which is lowly expressed in aggressive HER2 positive breast cancer, was also downregulated (20). We found *DUSP4* to be significantly downregulated, its decreased expression being associated with metastatic cancer (21). To have a detailed look between individual biological replicates, we plotted a heatmap of the top 30 differentially expressed genes in all samples, which further highlighted the distinct expression patterns between the two cell types (Fig. 2D). We identified a number of novel genes related to lapatinib-acquired drug resistance including *CSF3R, TLE6, GABRE,* and *PAD12*. Additionally, the low expression of *RAP1GAP*, a tumour suppressor gene, whose low expression levels is associated with epithelial-to-mesenchymal transition (EMT), a phenotype of aggressive cancers (22). These previously unknown genes could be further validated in physiologically relevant models and may serve as targets for lapatinib acquired drug resistance in HER2 positive breast cancer.

Our RNA-seq data revealed at least 17 differentially expressed histone-modifying enzymes that play a crucial role in the epigenetic regulation of key cellular functions and the post-translational modification of both histone and non-histone substrates (Fig. 2E). The findings showed significant (p value < 0.05) up-regulation of several genes encoding histone deacetylases including HDAC2, HDAC10 and several lysine demethylases (KDM) enzymes amongst others.

The pathway analysis indicated that the significantly differentially upregulated genes (DEGs) (adjusted p value <0.05) were mainly enriched in cell proliferation, angiogenesis and apoptotic process. This may imply that these biological processes enhance cell-to-cell contact, potentially leading to increased motility, indicating the ability to acquire new phenotypes for aggressive disease. Additionally, MAPK pathways, multicellular organism development and oxidation-reduction process indicate engagement of inflammatory responses, which could be a response to cellular stress (23).

### ATAC-seq identifies key regions driving lapatinib resistance in breast cancer cells

We used the ATAC-seq to profile the genome-wide accessibility landscape in SKBR3 and SKBR3-L cells. We isolated DNA from SKBR3-L cells and its corresponding lapatinib-sensitive parental cell lines in three biological replicates. Samples exhibited the expected periodicity of the insert length (supplementary fig. 1B and 1C). Hierarchical clustering based on the correlation of accessibility separated the samples into two groups: one containing the majority peaks of SKBR3 samples and the other containing the majority peaks of lapatinib-resistant samples, SKBR3-L (supplementary fig. 1D).

We identified 19,854 significantly differential peaks between SKBR3 and SKBR3-L cells with an adjusted p value of < 0.05. We conducted a comparison of differential accessible regions (DAR’s) between these two groups and discovered that SKBR3-L cells showed a decrease in chromatin accessibility with 12,591 peaks indicating lower inaccessibility, as the peaks had a fold-change < 0 with an adjusted p value < 0.05. 7,261 peaks showed accessibility with a fold-change > 0 and an adjusted p value < 0.05. We observed 1,213 regions as “hyper-accessible” (> 2-fold-change, adjusted p value < 0.05) and 1914 regions as “hypo-accessible” (< 2-fold-change, adjusted p value < 0.05) (Fig. 3A and supplementary fig. 1D). Through further quantitative comparison of the two cell types, we identified the resistant cell line to have greater inaccessibility with concentration changes at the higher end (≥ 8) reducing from 4,106 regions to 3,085 regions and the lower end (2-3.99) increasing from 911 regions to 1192 (supplementary fig. 2A).

**Figure 3.**
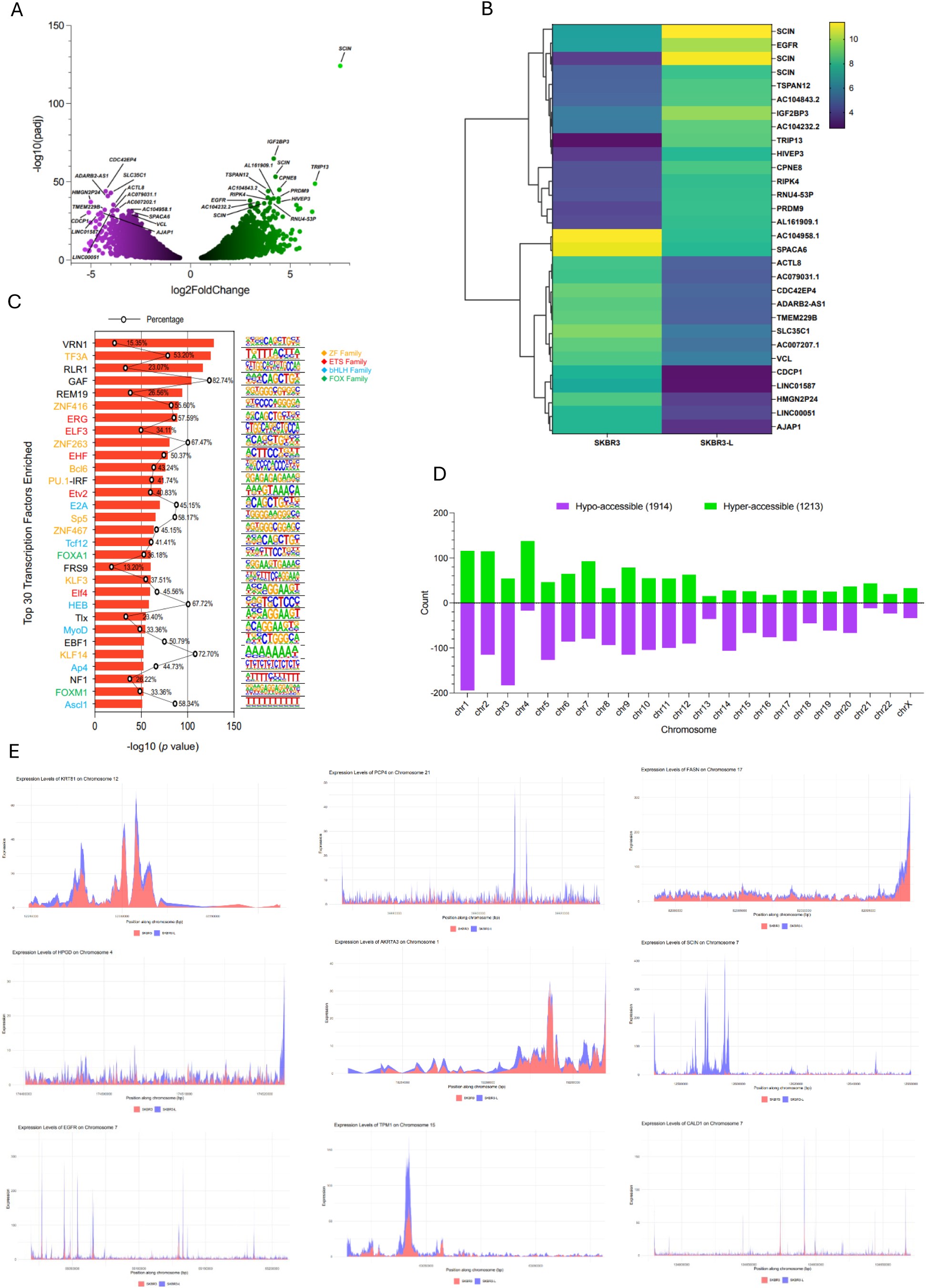
Chromatin accessibility changes between SKBR3 and SKBR3-L cell lines. **(A)** Volcano plot of differentially accessible regions with the top 30 significant regions labelled by gene, with log2fold change < -2, adjusted p value < 0.05 (purple). Upregulated regions are shown in red with log2fold change > 2, adjusted p value < 0.05 (green). **(B)** Hierarchical clustered heatmap of the top 30 significantly differentially expressed regions identified by gene. **(C)** Transcription factor enrichment motif analysis. Top 30 significant hits (adjusted p value < 1e-52) identified motifs and the transcriptional factors that bind to the region. **(D)** Comparison by chromosome of SKBR3 SKBR3-L cell lines. Hyper-accessible regions (log2fold change > 2, adjusted p value < 0.05) and hypo-accessible regions (log2fold change < -2, adjusted p value < 0.05) are plotted on the bar graph. **(E)** Candidate regions within 6bp from the TSS of nine different markers between SKBR3 and SKB3-L cells. SKBR3 (red), SKBR3-L (blue).

A hierarchical clustered heatmap was produced to visualise the expression profile of the top 30 DARs corresponding to the accessibility signal, sorted by their log2fold change of > 0.5, adjusted p value < 0.05 through plotting their log2 transformed expression change value (Fig. 3B). Through this we were able to identify and visualise co-regulated regions across treatment conditions. Through the linkage of known gene regions and chromatin states we confirmed *EGFR* to be highly accessible, known to give cells basal-like features (24). In addition, *SCIN,* a gene strongly associated with proliferation, migration and differentiation in cancer exhibited a with a (log2fold change of 7.52) high level of significance. *TRIP13* and *IGF2BP3*, which promotes tumour growth were also high differentially accessible (25,26).

Transcription factors (TFs) regulate gene expression by binding to specific DNA sequences in promoter regions (23). Chromatin accessibility is crucial for TF binding as it impacts the TFs ability to bind to the nucleosomes (24). Using ATAC-seq data and HOMER v.4.11 for motif enrichment analysis, we identified candidate TFs associated with genomic alterations by analysing differential chromatin accessibility in promoter or enhancer elements.

Our analysis identified 444 known motifs enriched with 27 de novo results that showed significant enrichment (adjusted p value < 0.05). We found ERG motif to be highly enriched alongside FOXM1, an oncogenic transcription factor known to be highly upregulated in breast cancer and is associated with poor patient survival, was significantly enriched in accessible regions (Fig. 3C) (27). Additionally, we identified several Zinc Finger (ZF) family members, which include the Kruppel-like factor (KLF) family members, to be significantly enriched, followed by others from Erythroblast Transformation Specific (ETS), basic Helix-Loop-Helix (bHLH), and Forkhead box (Fox). Each TF’s enrichment was quantified, with the most significant log10 > 50, adjusted p value < 0.05.

Furthermore, we categorised the differential peak analysis to each chromosome to assess the distribution of chromatin dynamics across the genome (Fig. 3D). The accessibility of the chromosomes vary with chromosome 4 showing the greatest increase in accessibility at 138 “hyper-accessible” regions identified. Aberrations in chromosome 4 are associated with several types of cancers for which 141 tumours secreted and 54 cancer associated proteins are encoded, revealing regions in chromosome 4 as a strong candidate for biomarker verification and drug target discovery research (28).

Furthermore, genomic annotation of differential accessible regions revealed a clear distribution pattern with 435 hyper-accessible regions predominantly located in gene-distal intergenic areas, these likely control the spatiotemporal and quantitative expression dynamics and activation of potential enhancer that affect transcriptional activity (supplementary fig. 2B). In contrast, 743 hypo-accessible regions were found within intron regions and aberrant intron retention has high association with several cancer types (29). Overall, we identified 3,888 regions with differential accessibility at or near the Transcription Start Site (TSS) (1kbp) of which 72.6% coincided with RNA seq data, confirming chromatin accessibility’s role in modulating gene expression.

Following differential peak analysis, the top 100 significant regions were extracted and filtered by TSS distance and those within 6 base pairs of the TSS had their chromatin landscape mapped. In total 15 out of the 100 significant regions were at or near the TSS. Genes identified included novel genes such as *HPGD, FASN and KRT81,* as well as *SCIN* and *TPM1* which have already been confirmed in acquired drug resistance mechanisms associated with HER2 positive breast cancer (Fig. 3E). These genomic regions may not only provide an understanding of how the chromatin accessibility landscape may contribute to different treatment response, but also aid in our understanding of how epigenetic/transcriptomic mechanisms may influence drug sensitivity and resistance (30,31). These regions could prove to be potential targets for future investigation for their role in drug resistance.

### Proteomic analysis confirmed markers identified in transcriptomic datasets

To interrogate the dynamic changes at the proteome level, we performed an unbiased proteomic analysis using liquid chromatography – tandem mass spectrometry (LC-MS/MS) between SKBR3 and SKBR3-L cells. In total, we identified 325 proteins with significant increased expression, log2fold change > 0.1, adjusted p value < 0.05, and 270 with significantly decreased expression, log2fold change < -0.1, adjusted p value < 0.05 (Fig. 4A). All proteins quantified in each cell type were visualised in a volcano plot depicting protein abundance changes between SKBR3 and SKBR3-L cells (Fig. 4B). The differentially expressed proteins were clustered and visualised as each biological replicate on a heatmap (Fig. 4C). Proteins within each cell type are clustered together and show consistent expression patterns.

**Figure 4.**
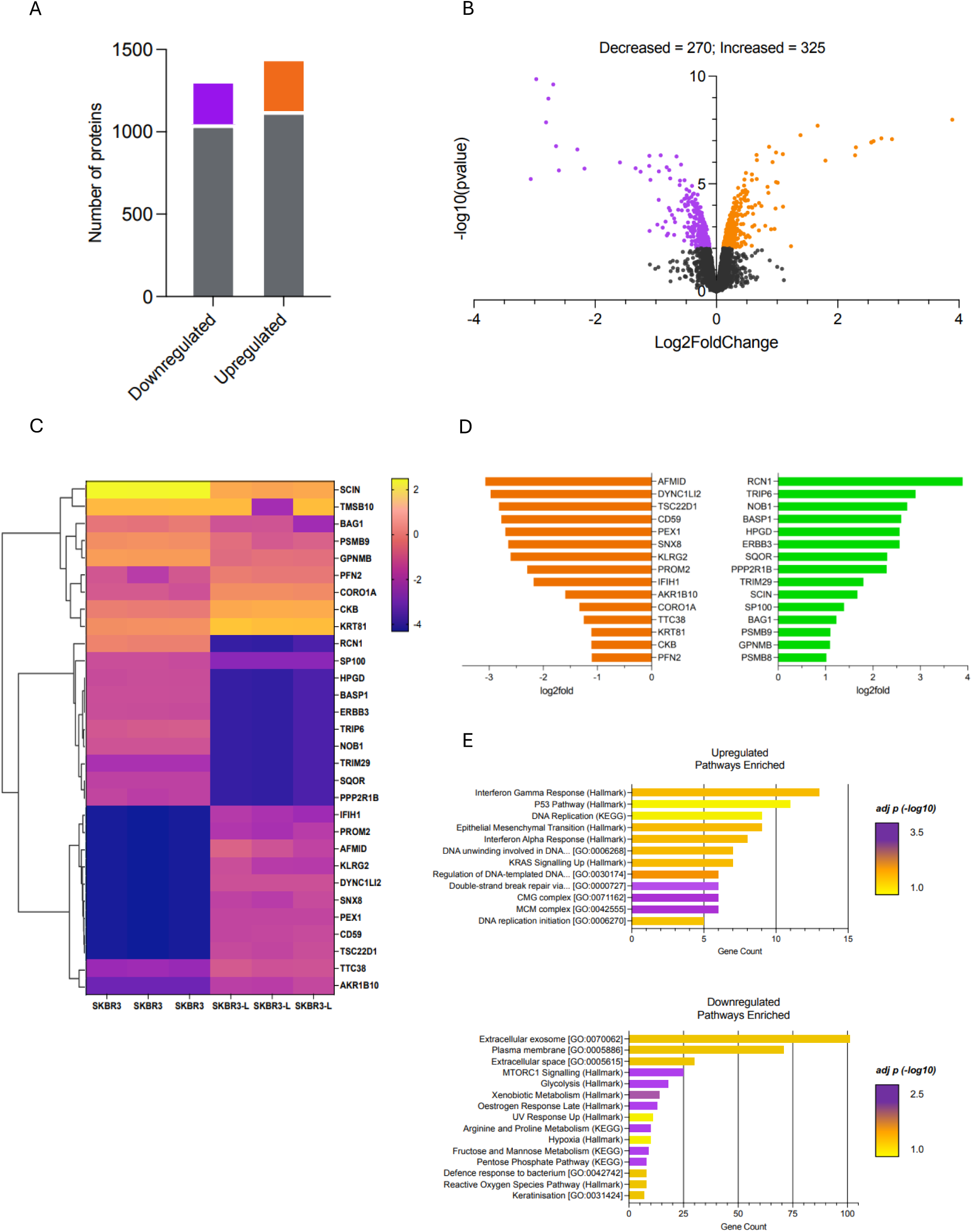
Proteomic analysis validated the markers found in transcriptomic datasets. **(A)** Number of differentially expressed proteins in SKBR3 compared with SKBR3-L cells. Statistical significance is shown as log2fold change >2, adjusted p-values < 0.05 for upregulated proteins and log2fold change < -2, adjusted p-values < 0.05 for downregulated proteins. **(B)** Volcano plot of differentially expressed proteins between SKBR3 and SKBR-L cells. **(C)** Heatmap of the top 30 significant differentially expressed genes labelled per each biological replicate. **(D)** Bar plot showing the top 15 downregulated and upregulated proteins based on their log2fold expression. **(E and F)** Bar plot showing biological pathways associated with significantly differentially expressed proteins (adjusted p value < 0.05).

We further visualised the protein expression change of the top 15 upregulated and top 15 downregulated proteins in a bar graph (Fig. 4D) for comparison. We found RCN1, TRIP6 and HPGD to be amongst those significantly upregulated, confirming some of the candidates we observed in our RNA-seq and ATAC-seq datasets. Whilst AFMID, CD59 and KRT81 to be significantly downregulated. PROM2 was amongst those downregulated and is of significant interest as its prognostic value in cancer is controversial, with its under expression potentially providing benefit to drug resistant cells (32).

Pathway enrichment analysis of both significantly (adjusted p value < 0.05) upregulated and downregulated proteins confirmed pathways identified in RNA seq (Fig. 2F) were also identified in proteomic (Fig. 4E). Upregulated pathways showed enrichment of proteins implicated in Epithelial Mesenchymal Transition (EMT) process. Additionally, double strand break repair was also enriched, which contributes to genomic instability. KRAS signalling, which is highly intertwined with HER2 signalling, plays a key role in cell survival, proliferation and migration. Downregulated pathways included extracellular exosome – small extracellular vesicles that play a role in cell-to-cell communication and mTORC1, which is involved in signal integration from multiple growth factors and energy supplies to promote cell growth (33).

### A combined approach to unravelling resistance culminates in 9 candidate markers

To unravel the complex interactions and resistance mechanisms between SKBR3 and SKBR-L cells, we employed an integrated multi-omic approach, combining ATAC-seq, RNA-seq, and proteomics data. This comprehensive data analysis led to the identification of 83 significant differential changes (showcased in a Venn diagram) revealing shared genomic regions that could be key targets for future research (Fig. 5A). We further dissected these interactions using volcano plots, providing a detailed view of the relationships between differentially expressed genes, accessible chromatin regions, and proteins (Fig. 5B).

**Figure 5.**
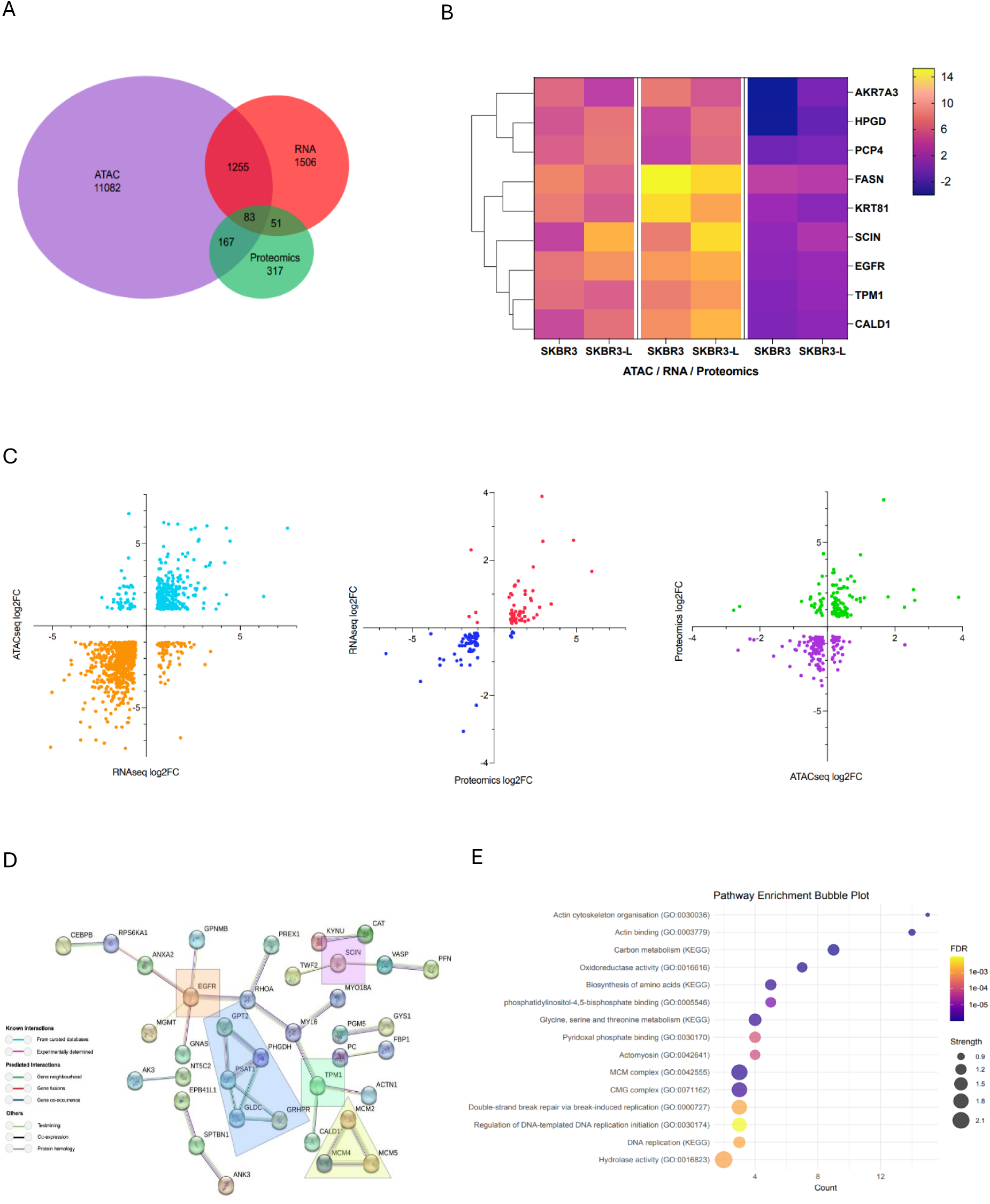
A combined approach to investigating lapatinib resistance resulted in the identification of 9 candidate markers. **(A)** Overlapping regions between ATAC-seq (log fold change > 1, adjusted p value < 0.05), RNA seq (log fold change > 1, adjusted p value < 0.05 and) and proteomics (log fold change > 0.1, adjusted p value < 0.05 and). **(B)** Heatmap of 9 candidate genes identified as differentially expressed/accessible across the 3 datasets. **(C)** Volcano plots of significantly differentially regulated interactions showing RNA-seq vs ATAC-seq, proteomics vs RNA-seq, ATAC-seq vs proteomics. **(F)** Protein-protein interaction network. The markers encoded by the common 83 identified genes as identified in the 3 datasets. Analysed with String v.12.0 and mapped; nodes indicating proteins and different coloured edges representing protein-protein associations amongst neighbouring nodes. **(G)** Biological pathways associated with the 83 significant differentially expressed genes as observed in the 3 datasets.

Focusing on key candidate markers in the three datasets, we narrowed down our list to 9 markers with significant changes across ATAC-seq, RNA-seq, and proteomic data, suggesting their potential roles in the development of lapatinib resistance. These markers were visualised using a hierarchical clustered heatmap, which displayed the co-regulated expression patterns across different treatment conditions in the three datasets (Fig. 5C). Notably, genes such as *SCIN* and *EGFR*, already known to be involved in HER2-positive breast cancer were confirmed, reinforcing the validity of our approach. Other significant novel candidate genes include *HPGD*, *TPM1*, *CALD1*, *PCP4*, *AKR7A3*, *KRT81*, and *FASN*, which are anticipated to be pivotal in lapatinib resistant mechanisms.

To reveal the protein-protein interaction (PPI) network and co-expression of the differential expressed proteins identified in our integrated analysis (adjusted p value < 0.05), we utilised String v.12.0 to analyse known and predicted physical interactions and functional associations between proteins (Fig. 5D). We used a high co-expression/interaction base score (0.700) to ensure high accuracy, and our PPI enrichment returned an overall p value of 0.0000137 with an average local cluster coefficient of 0.327. Proteins such as EGFR and TPM1 are centrally located and have multiple connections displayed within the network suggesting their key role in driving lapatinib acquired drug resistance.

A pathway enrichment analysis of the 83 significant proteins (Fig. 5E) revealed significant enrichment (adjusted p value < 0.05) in pathways related to actin remodelling, carbon metabolism, and double strand break repair. Actin remodelling influences tumour growth and chemoresistance, with aberrant actin isoforms serving as cancer biomarkers (34). Carbon metabolism plays a role in tumorigenesis and drug resistance via DNA methylation, while impaired double strand break repair contributes to genomic instability (35).

## Discussion

Acquired drug resistance is a high probability event in breast cancer in which cancer cells must remodel and rewire their genome, epigenome and proteome to be able to evade treatment and molecularly evolve into an incurable disease. Despite advancements in targeted therapies like lapatinib, drug resistance in HER2-positive breast cancer persists with over 70% of patients relapsing after initial treatment (1). This high frequency of acquired drug resistance in breast cancer cells could highlight their inherent ability to relapse. In contrast, our work shows that HER2 positive breast cancer cells gain the ability to become resistant to lapatinib through the dramatic remodelling of the chromatin, transcriptome and the proteome (Fig. 5B and 5C). Surprisingly, our data demonstrate that the aggressive phenotype observed in drug resistance correlates with downregulation in gene expression patterns and condensed chromatin landscape associated with limited, yet robust changes in the proteome. Our findings showcase the strength of integrating genome-wide molecular techniques using a genetic model system to reveal previously unnoticed variations in cellular states across multiple levels of regulation.

Across many cancer types, increased chromatin accessibility (36,37) and gene expression changes correlate with aggressiveness and poor patient outcomes (38). However, in our *in vitro* model, the link between increased chromatin accessibility and aggressiveness was not observed, as our investigation revealed a predominant decrease in chromatin peaks associated with significant downregulation of gene expression, suggesting a reduction of accessibility was associated with resistance mechanisms. Our data indicates that when developing lapatinib-acquired drug resistance, cells establish an invasion promoting gene expression programme through stabilising chromatin dynamics at specific regulatory genes. AKR7A3, a member of the aldo-keto reductase (AKR) protein family is downregulated in SKBR3-L cells (Fig. 3E and 5B). Its downregulation is associated with metastasis in pancreatic ductal carcinoma (39). We observed elevated expression in other key regions, such as EGFR and SCIN, which are well-known to be highly expressed in breast cancer resistance. We saw significant transcriptional changes including the downregulation of key markers like EGR1 and CD24, whose reduced expression is linked to increased invasiveness and drug resistance and identified proteins such as TRIP6 and TRIM29 that enhance (cancer stem cells) CSC-like properties (40,41). Our analysis identified nine candidates (Fig. 3E, 5B, and 5C) that were significantly differentially expressed across all three datasets, including seven novel candidates, as being involved in drug resistance in HER2 positive breast cancer. We think that, despite the predominant chromatin closing and the corresponding downregulation of gene expression with minimal proteome changes, certain key regulatory elements undergo significant shifts during acquired drug resistance. These changes likely drive the phenotypic alterations in drug resistance observed in our *in vitro* model. Chromatin closing results in the downregulation of genes linked to disruptive changes in cell morphology and motility (Fig 1C, 1D, 1E). Meanwhile, other regions with differentially accessible chromatin suggest network rewiring, contributing to the development of an aggressive phenotype during acquired drug resistance.

Our data suggests that SKBR3-L cells exist in a “pre-emptive” chromatin configuration state that has an overall inaccessible chromatin dynamic, which still permits the binding of specific motifs during drug resistance. Our results show the enrichment of several transcription factors, particularly pioneer factors such as Foxa, KLF and Ascl1 (Fig. 3C). These enriched motifs could establish this “pre-emptive” architecture that bind their target sites in closed chromatin and subsequent chromatin opening (Fig. 3E and 5C) (42). This could explain the pattern of overall chromatin closing in SKBR3-L cells being observed concurrently with enhanced accessibility near the transcriptional start site, particular within 6 base pairs of critical genes involved in drug resistance.

The aggressive nature of lapatinib-resistant cells is evident compared to its cancer counterpart, as our analysis shows significant anchorage-independent growth (Fig. 1D), which correlates with hyperactivation of the MAPK and KRAS pathways (Fig. 2F and 4E), leading to an enhanced transformational potential (Fig. 1D and 1E) (43). Upregulation of upstream receptors, such as HER2 and EGFR – key activators of the MAPK pathway, also contributes to enhanced growth and drug resistance (44–46). Additionally, morphological changes, including loss of sphericity and increased overall cell size (Fig. 1C), suggest a loss of cell-to-cell contact and communication, indicative of invasive behaviour. This is consistent with our protein-protein interaction analysis, where pathways related to cytoskeletal organisation and cell adhesion were highly enriched (Fig. 5F). These alterations are often associated with a transition to a mesenchymal phenotype (Fig. 4E), which is characterised by increased motility, invasiveness, and drug resistance, which are features of metastatic state (47).

While the gene expression patterns of lapatinib-sensitive cells at the baseline level displayed some diversity between biological replicates, cells that developed drug resistance exhibited a notable convergence in their expression profiles (Fig. 2A). This uniformity in adaptive responses among the resistant cells is potentially due to the selective pressure of lapatinib with resistant cells activating similar pathways that lead to a stabilisation of specific gene expression patterns. A significant proportion (∼ 46%) of transcriptional changes coincided with significant chromatin alternations (Fig. 5A). Alongside this, a notable overall decrease in gene expression patterns correlated with genes showing closed chromatin states at the promoter region in the resistant cell line, underscoring the epigenome’s role as a regulator of resistance in HER2-positive breast cancer. From an epigenetic regulation standpoint, we examined changes in the expression of known epigenetic enzymes (Fig. 2E). The findings indicate that the upregulation of HDACs may play a role in the downregulation of specific genes associated with lapatinib resistance. Previous studies have explored the epigenomic context of trastuzumab resistance with hypermethylation of MCF7 cells being identified as a driver of resistance; nonetheless, few studies have emerged on SKBR3 cell lines which are associated with a more invasive and aggressive phenotype (48,49).

We took a target discovery approach using a combined molecular analysis, which led to the identification of lapatinib acquired drug resistance signature (Fig. 5B), serving as a potential target for reversing drug resistance. These target genes and their associated pathways not only shed light on the mechanistic underpinnings of drug resistance, but also highlight the complex interplay between genetic and epigenetic factors in cancer progression. The use of multi-omics approach has recently emerged as a promising technique to identify previously unknown drug targets. Given the dramatic remodelling of the key gene regulatory state as a result of changes in the chromatin, which leads to changes in gene expression and proteins, we speculate that other cancer types may also have similar signature as we have observed. Our target discovery approach provides the scientific community future research incentive to study this signature in models that are physiologically relevant to the human context, and if confirmed, to target it with inhibitors and gene manipulation technologies as a basis for potential biomarkers in acquired drug resistance.

## Supplementary data

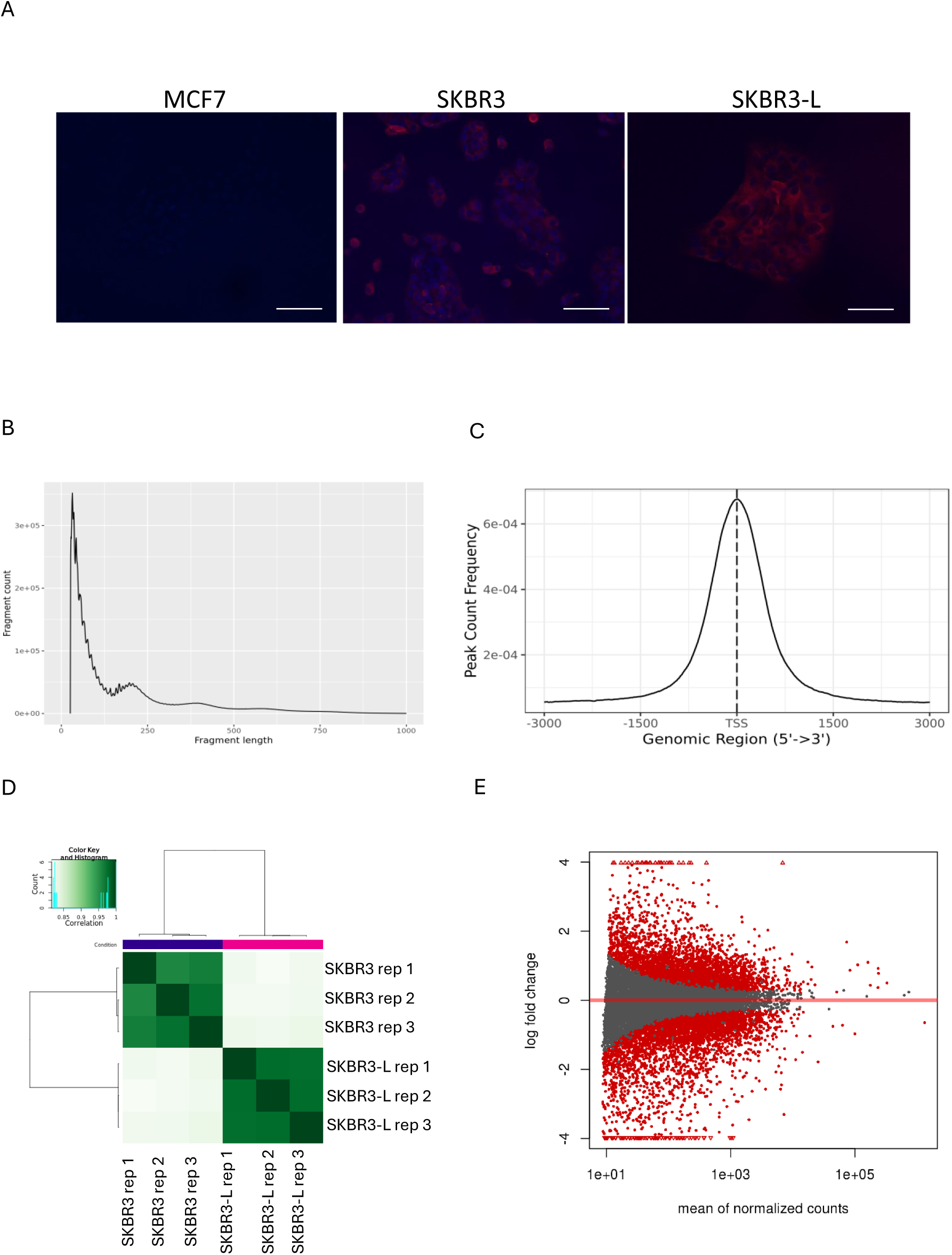
**(A)** Additional fluorescence microscopy images confirming the expression of HER2 protein in both cell types, with MCF7 cells used as negative controls, which do not express the HER2 protein. 20x magnification. Scale bars: 100μm. **(B)** The representative insert size distribution demonstrates clear modulation of the signal, with distinct peaks corresponding to both mono-nucleosomes and di-nucleosomes, indicating well-defined nucleosome positioning. **(C)** Representative plot of aggregate signal around transcription start sites (TSS) shows strong enrichment of reads near TSS regions. **(D)** Correlation of chromatin accessibility between different cell lines and across technical replicates. **(E)** MA Plot of differential accessibility (log2 fold change in reads per accessible region) versus the mean number of reads per region, adjusted p value <0.05.

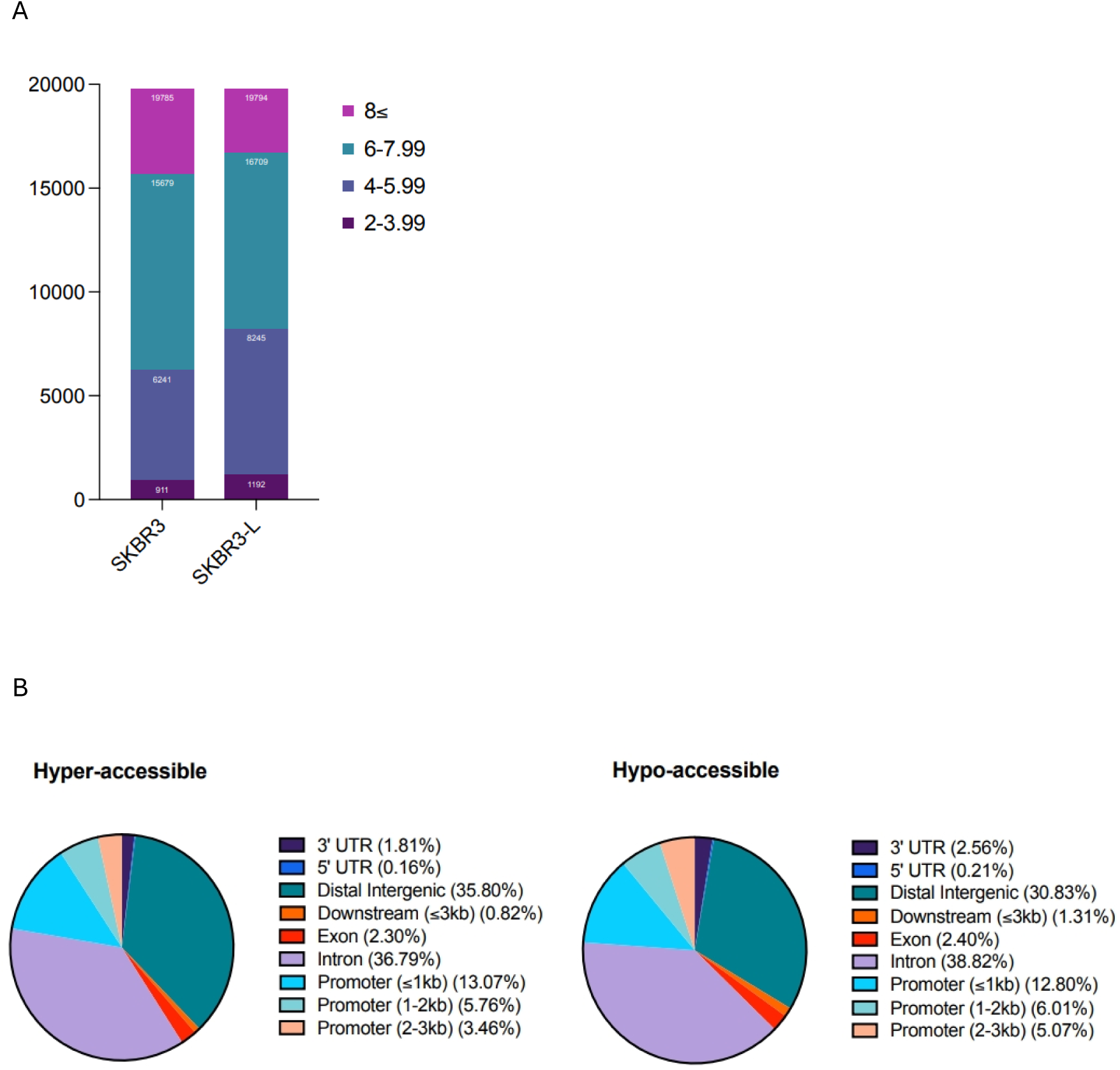
**(A)** A comparison of SKBR3 SKBR3-L cell lines based on their significant differential logfold changes with adjusted p value of < 0.05. **(B)** Distance to the nearest transcription start sites (TSSs) for all differentially accessible regions in SKBR3 and SKBR3-L cells (p value of <0.05).

## Methods

### Cell culture

The SKBR3 and SKBR3-L cell lines were gifted by (6) and grown under recommended conditions. Briefly, SKBR3 and SKBR3-L cell medium consists of McCoy’s 5A Medium (ThermoFisher, #16600082) supplemented with 5% horse serum (Sigma-Aldrich, #H1138) and 1× penicillin/streptomycin (Pen/Strep). The SKBR3-L cells were cultured in 2μM Lapatinib (BioTechne, #6811/10).

SKBR3 cells were cultured in increasing concentrations of lapatinib, starting approximately from the IC50 value, respectively 77nM, as determined by dose-response studies. Through continuous exposure to the drug, the lapatinib sensitive cells die and the live cells are cultured until they outgrow the lapatinib dose. The dose is increased, and the process is repeated until the cells are able to grow in 2μM of lapatinib, a concentration equal to twice the resistance threshold.

### 3D Cell Culture and morphology analysis

3D cell culture was performed using pre-made BIO-FLOAT 96-well plates (Sarstedt, #83.3925.400) with a round base. 1000 cells were plated onto the 96-well plates and cells/spheroids were immediately visualised and tracked using LiveCyte2 quantitative phase imaging (QPI) for 12 hours to characterise the changes in morphology. Morphological changes were analysed using software (Acquire) embedded in the microscope and metrics such as size, area and motility were processed.

### Soft Agar Colony Formation Assay

A 0.8% base layer mixed with cell medium was formed in plates using ultra-pure culture grade agarose (Thermo Fisher Scientific, #16500500), allowed to solidify at room temperature for 30 minutes. 5000 cells per well were mixed with 0.35% ultra-pure agarose with cell medium and plated evenly, drop wise, on top of the bottom layer for overlay culture. Medium was changed every 3 days for 3 weeks. Colonies were fixed using 4% PFA and permeabilised using 100% methanol. Colonies were stained using 0.05% Crystal Violet dye (APC Pure, #RRSP23-D), and images were taken using a brightfield microscope (Nikon Ni-E, fitted with the Nikon DS-Fi3 camera) capturing most of the perimeter of the well. Binary masks were applied to each of the images, and thresholding parameters for diameter ranging from 30μm to 50μm were set on ImageJ. Colonies were counted using ImageJ only if they satisfied criteria above the threshold values, and colony counts were then manually checked and adjusted if necessary.

### RNA Extraction, Library Preparation and NovaSeq Sequencing

Total RNA was extracted from frozen cell pellets using Qiagen RNeasy Mini kit following manufacturer’s instructions (Qiagen, Hilden, Germany). RNA samples were quantified using Qubit 4.0 Fluorometer (Life Technologies, Carlsbad, CA, USA) and RNA integrity was checked with RNA Kit on Agilent 5300 Fragment Analyser (Agilent Technologies, Palo Alto, CA, USA). RNA sequencing libraries were prepared using the NEBNext Ultra RNA Library Prep Kit for Illumina following manufacturer’s instructions (NEB #E7770, Ipswich, MA, USA). Briefly, mRNAs were first enriched with Oligo(dT) beads. Enriched mRNAs were fragmented for 15 minutes at 94 °C. First strand and second strand cDNAs were subsequently synthesised. cDNA fragments were end repaired and adenylated at 3’ends, and universal adapters were ligated to cDNA fragments, followed by index addition and library enrichment by limited-cycle PCR. Sequencing libraries were validated using NGS Kit on the Agilent 5300 Fragment Analyser (Agilent Technologies, Palo Alto, CA, USA), and quantified by using Qubit 4.0 Fluorometer (Invitrogen, Carlsbad, CA). The sequencing libraries were multiplexed and loaded on the flowcell on the Illumina NovaSeq instrument according to manufacturer’s instructions. The samples were sequenced using a 2×150 Pair-End (PE) configuration v1.5. Image analysis and base calling were conducted by the NovaSeq Control Software v1.7 on the NovaSeq instrument. Raw sequence data (.bcl files) generated from Illumina NovaSeq was converted into fastq files and de-multiplexed using Illumina bcl2fastq program version 2.20. One mismatch was allowed for index sequence identification.

### Data analysis of RNA-seq

After investigating the quality of the raw data, sequence reads were trimmed to remove possible adapter sequences and nucleotides with poor quality using Trimmomatic v.0.36. The trimmed reads were mapped to the Homo sapiens reference genome available on ENSEMBL using the STAR aligner v.2.5.2b. The STAR aligner is a splice aligner that detects splice junctions and incorporates them to help align the entire read sequences to generate BAM files. Unique gene hit counts were calculated by using feature Counts from the Subread package v.1.5.2. Only unique reads that fell within exon regions were counted. After extraction of gene hit counts, the gene hit counts table was used for downstream differential expression analysis. Using DESeq2, a comparison of gene expression between the groups of samples was performed. The Wald test was used to generate p-values and Log2 fold changes. Genes with adjusted p values ≤ 0.05 and absolute log2 fold changes ≥ 1 were identified as differentially expressed genes for each comparison. A gene ontology analysis was performed on the statistically significant set of genes by implementing the software GeneSCF v.1.1 (Subhash and kanduri, 2016). The goa_human GO list was used to cluster the set of genes based on their biological process and determine their statistical significance. A PCA analysis was performed using the “plotPCA” function within the DESeq2 R package (Love *et al.,* 2014). The plot was created in Graphpad Prism v.10.0.3 (GraphPad Software, 2023) and shows the samples in a 2D plane spanned by their first two principal components. The top 500 genes, selected by highest row variance, were used to generate the plot. Subsequent plots and graphs were produced using R v.4.4.0 (R Core Team, 2023) and GraphPad Prism v.10.0.3 (GraphPad Software, 2023).

### ATAC-seq Library Preparation and Sequencing

Live cell samples were quantified and assessed for viability using a Countess Automated Cell Counter (ThermoFisher Scientific, Waltham, MA, USA). After cell lysis and cytosol removal, nuclei were treated with Tn5 enzyme (Illumina, #20034197) for 30 minutes at 37°C and purified with Minelute PCR Purification Kit (Qiagen, #28004) to produce tagmented DNA samples. Tagmented DNA was barcoded with Nextera Index Kit v2 (Illumina, #FC-131-2001) and amplified via PCR prior to a SPRI Bead cleanup to yield purified DNA libraries. The sequencing libraries were clustered on a NovaSeq flowcell. After clustering, the flowcell was loaded on the Illumina NovaSeq instrument according to manufacturer’s instructions. The samples were sequenced using a 2×150bp Paired End (PE) configuration. Image analysis and base calling were conducted by the Control Software (CS). Raw sequence data (.bcl files) generated from Illumina instrument was converted into fastq files and de-multiplexed using Illumina’s bcl2fastq 2.17 software. One mismatch was allowed for index sequence identification.

### Data analysis of ATAC-seq

After investigating the quality of the raw data, sequencing adapters and low-quality bases were trimmed using Trimmomatic 0.38. Cleaned reads were then aligned to reference genome GRCm38 using bowtie2. Aligned reads were filtered using samtools 1.9 to keep alignments that (1) have a minimum mapping quality of 30, (2) were aligned concordantly, and (3) were the primary called alignments. PCR or optical duplicates were marked using Picard v.2.18.26 and removed. Prior to peak calling, reads mapping to mitochondria (mt) were called and filtered, and reads mapping to unplaced contigs were removed. MACS2 v.2.1.2 (Zhang *et al.,* 2008) was used for peak calling to identify open chromatin regions. Valid peaks from each group or condition were then merged and peaks called in at least 66% of samples were kept for downstream analyses. For each pair-wise comparison, peaks from SKBR3 and SKBR3-L were merged, and peaks found in either condition were kept for downstream analyses. Reads falling beneath peaks were counted in all samples, and these counts were used for differential peak analyses using the R package DiffBind v.3.14 (Stark and Brown, 2011). BED files were then loaded into HOMER v.5.0 for MOTIF enrichment analysis and the top 30 known transcription factors identified. Plots and graphs were produced using R v.4.4.0 (R Core Team, 2023) and GraphPad Prism v.10.0.3 (GraphPad Software, 2023).

### Proteomic sample preparation

For proteomic experiments, cells were grown in 2D cell cultures. 30μg of protein were subjected to cysteine reduction and alkylation using sequential incubation with 10 mM dithiothreitol and 16.6 mM iodoacetamide for 1 h and 30 min, respectively, at 25 °C with agitation. Trypsin beads (50% slurry of TLCK-trypsin) were equilibrated with three washes with 20 mM HEPES (pH 8.0), the urea concentration in the protein suspensions was reduced to 2 M by the addition of 600 µL of 20 mM HEPES (pH 8.0), 70 μL of equilibrated trypsin beads were added and samples were incubated overnight at 37 °C. Trypsin beads were removed by centrifugation (2000 × g at 5 °C for 5 min).

Peptide solutions were desalted using the AssayMAP Bravo (Agilent Technologies) platform. For desalting, protocol peptide clean-up v3.0 was used. Reverse phase S cartridges (Agilent, 5 μL bed volume) were primed with 250 μL 99.9% acetonitrile (ACN) with 0.1%TFA and equilibrated with 250 μL 0.1% TFA at a flow rate of 10 μL/min. The samples were loaded at 20 μL/min, followed by an internal cartridge wash with 0.1% TFA at a flow rate of 10 μL/min. Peptides were then eluted with 105 μL of solution (70/30 ACN/ H2O + 0.1% TFA. Eluted peptide solutions were dried in a SpeedVac vacuum concentrator and peptide pellets were stored at −80 °C. Samples were processed and run in the mass spectrometry in consecutive days in order to reduce technical variability. Peptide pellets were reconstituted in 5 µL of 0.1% TFA, 2 µL of this solution were further diluted in 18 µL of in 0.1% TFA and 2 µL were injected into the LC-MS/MS system. The LC-MS/MS platform consisted of a Dionex Ultimate 3000 RSLC coupled to Q Exactive™ Plus Orbitrap Mass Spectrometer (Thermo Fisher Scientific) through an EASY-Spray source. Mobile phases for the chromatographic separation of the peptides consisted of Solvent A (3% ACN; 0.1% FA) and Solvent B (99.9% ACN; 0.1% FA). Peptides were loaded in a μ-pre-column and separated in an analytical column using a gradient running from 3 to 28% B over 90 min. The UPLC system delivered a flow of 10 µL/min (loading) and 250 nL/min (gradient elution). The Q Exactive Plus operated a duty cycle of 2.1 s. Thus, it acquired full scan survey spectra (m/z 375–1500) with a 70,000 FWHM resolution followed by data-dependent acquisition in which the 15 most intense ions were selected for HCD (higher energy collisional dissociation) and MS/MS scanning (200–2000 m/z) with a resolution of 17,500 FWHM. A dynamic exclusion period of 30 s was enabled with m/z window of ±10 ppm. Mass spectrometry data collection was carried out using Thermo Scientific FreeStyle 1.4.

### Data analysis for proteomics

MS raw files were converted into mzML using MSConvert, part of the ProteoWizard software package (5). FragPipe (version v22.0) (6) with MSFragger (version 4.1) (7), Percolator (3.07.01) (8) and PTM-Shepherd (v2.0.5) (9) to analyze the data. Peptide and phosphopeptide quantification were performed using in-house software Pescal as described before (3). The resulting quantitative data was further analysed using Protools 2 R package(https://github.com/CutillasLab/protools2/releases/tag/v0.2.12.)

### Statistical Analysis

R v.4.4.0 (R Core Team, 2023) [programming language] and GraphPad Prism v.10.0.3 (GraphPad Software, 2023) were used for statistical analysis. This included data refinement, transformations, normalisations, and descriptive statistics. Alongside this R v.4.4.0 (R Core Team, 2023) and GraphPad Prism v.10.0.3 (GraphPad Software, 2023) were also used to illustrate our findings utilising an array of heatmaps, dendrograms, volcano plots, bubble plots and bar graphs. Experiments were all carried out in triplicates and the False Discovery Rate was used to control type 1 error ensuring high accuracy in our findings.

## Acknowledgements

We thank Dr Salvatore Federico Pedicona for critically reading the manuscript. We thanks the City, St George’s University of London’s imaging research facility (IRF) for the shared microscopy services.

## Authors contribution

Conceptualisation: A.H.; Methodology: A.H., J.S., V.R; Software: A.H., J.S., V.R., N.E.A; Validation: A.H., J.S.; Formal analysis: A.H., J.S., V.R., N.E.A; Resources: A.H., J.S.; Data curation: J.S. A.H., V.R., N.E.A; Writing - original draft: A.H., J.S.; Writing - review & editing: A.H., J.S.; Visualization: A.H., J.S.; Supervision: A.H.; Project administration: A.H.; Funding acquisition: A.H.

## Data availability

The BioProject ID for the RNA-seq and ATAC-seq data reported in this paper is: PRJNA1170334. The mass spectrometry-based proteomics data have been deposited to the ProteomeXchange Consortium via the PRIDE Project accession: PXD056848.

